# Improved Deep Learning Prediction of TCR-HLA Associations

**DOI:** 10.1101/2024.11.22.624910

**Authors:** Fumin Li, Si Liu, Wei Sun

## Abstract

Understanding the relationship between T cell receptors (TCRs) and human leukocyte antigens (HLAs) is essential for elucidating immune response specificity, uncovering mechanisms of autoimmunity, and advancing targeted immunotherapies. We have previously developed a deep learning method, DePTH (Deep Learning Prediction of TCR-HLA associations), to predict the association between a TCR and an HLA based on their amino acid sequences. In this work, we evaluated the performance of DePTH in two additional datasets, and investigated the influence of two potential confounding factors: TCR generation probability and the sequence length of CDR3 (Complementarity-Determining Region 3), which is a key region in the antigen-binding site of TCRs. Building on these insights, we combined training data from two datasets to train a new version of DePTH: DePTH 2.0.

## Introduction

Human Leukocyte Antigen (HLA) proteins and T cell receptors are two critical components of human immune system. There are tens of thousands of HLA alleles within the human population, each encoding a protein that presents peptides on cell surfaces to be examined by T cells. When T cells detect foreign antigens (e.g., peptides from virus), they initiate an immune response to eliminate cells presenting these antigens. This recognition process relies on T cell receptors (TCRs). A healthy individual possesses a repertoire of tens of millions of TCRs [6]. Most of these TCRs are rare since they are generated by a stochastic process with a huge number of possibilities (e.g., ∼ 10^14^ choices for TCR beta chains [3]). However, some TCRs are shared among multiple individuals and they are referred to as public TCRs. Public TCRs arise for two primary reasons. First, a TCR may be shared because it recognizes a common antigen, e.g., an antigen from common cold virus. Second, some TCRs have higher probabilities being generated [11].

By analyzing co-occurrence patterns of HLA alleles and public TCRs within human populations, we can identify associations between specific HLA alleles and TCRs. Previously, we leveraged these co-occurred TCR-HLA pairs to train a neural network named DePTH (Deep learning Prediction of TCR-HLA association) to predict TCR-HLA associations based on their amino acid sequences [9]. DePTH was trained using the HLA and TCR data of 666 individuals from Emerson et al. [6].

In this work, we evaluate DePTH’s performance using two new datasets: the Delmonte et al. dataset [5] and the TCGA dataset [16], which contain TCR and HLA data from 540 and 9,742 individuals, respectively. Our findings demonstrate that DePTH model is generalizable and it performs well in these two new datasets. Next, we study the impact of two factors that are associated with DePTH score: the TCR generation probability (TGP) and the length of the CDR3 sequence, which is the key section of TCR sequence that interacts with HLAs (Figure 1). TGP is negatively associated with DePTH score and CDR3 length is positively associated with DePTH score. We have also shown current training data selection procedure makes DePTH’s performance robust to these factors. Finally, using the additional data from Delmonte dataset [5], we train a new version of DePTH: DePTH 2.0.

**Figure 1:**
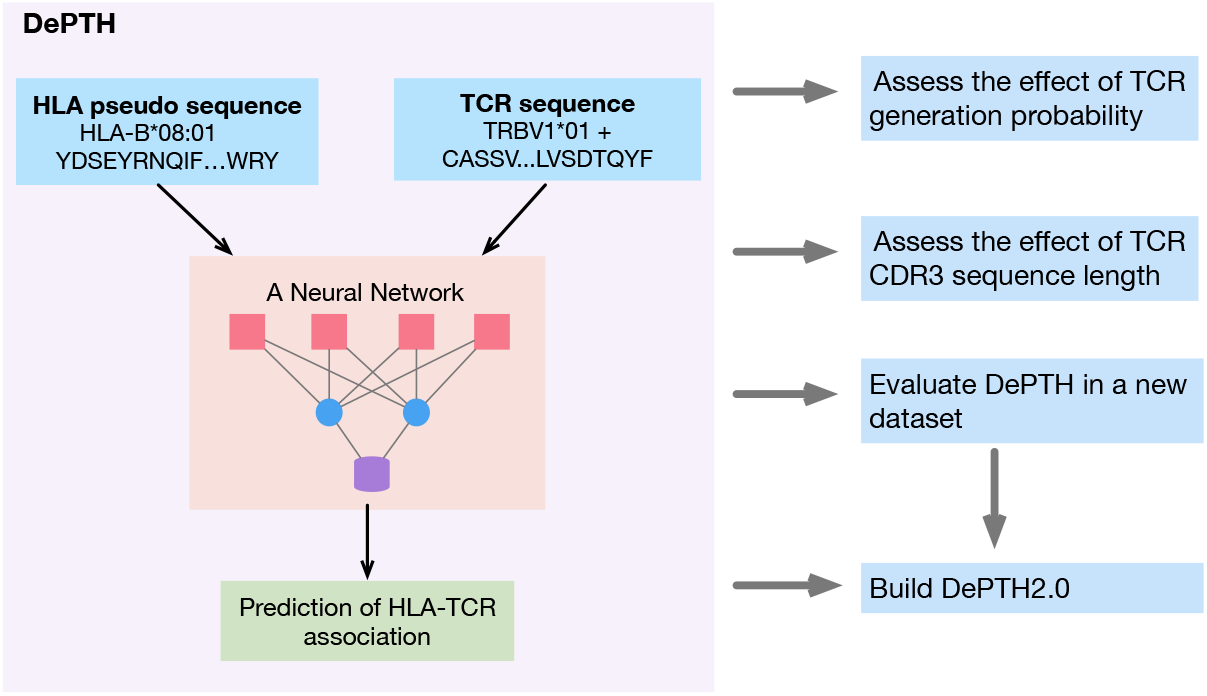
An overview of the DePTH model and evaluation/extension of DePTH in this paper.

## Methods

### A brief review of TCR, HLA and DePTH

The vast majority of T cells (∼ 95%) express alpha-beta TCRs that are composed of an alpha chain and a beta chain [1]. Each chain comprises a variable region, which is responsible for antigen recognition, and a constant region, which anchors a TCR to T-cell membrane. Within the variable region, three complementarity-determining regions (CDRs)—CDR1, CDR2, and CDR3—play a direct role in antigen interaction. CDR1 and CDR2 are encoded by the germline copy of the V (Variable) gene segment. In contrast, CDR3 is generated through stochastic recombination of the V, D (Diversity, used by beta but not alpha chain), and J (Joining) gene segments, followed by additional somatic mutations [8]. This process gives rise to the immense diversity of the TCR repertoire. A healthy individual may have 10^7^ or more unique TCR beta chains [6]. Most TCR data from bulk tissue samples, including the Emerson dataset [6] used to train our DePTH model, contain only the data pertaining to the beta chain. Therefore DePTH and the newly developed DePTH are limited to assess TCR beta chains.

HLA represents the human version of the Major Histocompatibility Complex (MHC). The genomic loci encoding HLA are among the most polymorphic regions in the human genome, with over 15,000 common HLA alleles documented [4, 13, 18]. An individual can possess up to 16 different HLA alleles, including six HLA class I alleles—comprising two HLA-A, two HLA-B, and two HLA-C alleles, and ten HLA class II alleles, which include two HLA-DR, four HLA-DP, and four HLA-DQ alleles.

In the immune system, peptides are presented through the peptide-binding groove of HLA molecules. A TCR engages these peptides through its CDR regions, establishing spatial complementarity with the HLA-peptide complex. This interaction triggers a cascade of signal transduction events within the T cell, ultimately leading to T cell activation. Activated T cells then secrete cytokines, proliferate, and differentiate, driving the immune response.

It is well-known that TCRs’ capabilities to recognize antigens can be restricted to certain HLA alleles. A method that can quantify such HLA restrictions can be useful for many studies of immune function. Towards this end, we developed DePTH, a neural network that uses the amino acid sequences of both TCRs and HLAs to predict the association between a TCR and an HLA allele (Figure 1) [9]. Instead of treating HLA allele as a categorical variable, we represent an HLA allele based on its sequence. This strategy allows DePTH to be applicable for any HLA alleles with sequence information and effectively borrow information across HLA alleles with similar sequences.

### Two additional datasets: Delmonte and TCGA

We previously evaluated the performance of DePTH using the Emerson dataset [6] and several other smaller datasets with TCR-HLA interaction annotation [9, 17, 15]. However, DePTH’s generalizability across larger and more diverse datasets remains unexplored. In this work, we evaluated the performance of the DePTH on two large datasets: the Delmonte dataset [5] and the TCGA dataset [16].

The Delmonte dataset [5] includes the HLA and TCR beta chain data from 540 COVID-19 patients. The TCR sequencing depth in Delmonte is comparable to that of the Emerson dataset (Supplementary Figure 6). When focusing on TCR-HLA pairs where both the HLA and the TCR appear in at least five individuals, the dataset comprises approximately 100 million TCR-HLA pairs for HLA-I alleles and 175 million pairs for HLA-II alleles. Among these 880 pairs were identified as positive (i.e., significant TCR-HLA associations) for HLA-I and 4,505 pairs for HLA-II, with a false discovery rate (FDR) *<* 0.1.

The TCGA dataset [16] differs significantly from the Emerson and Delmonte datasets in two key aspects. First, the TCR data were extracted from bulk RNA-seq, instead of TCR-seq, from tumor tissues. Limited amount of T cell infiltration in tumor and limited sequence coverage on TCR genes both reduce the amount of TCR sequences per sample. Second, while individual samples contain limited number of TCRs, the sample size is large, with bulk RNA-seq data from 9,736 cancer patients. In total, the TCGA dataset includes 211,346 TCR beta chain records. However, it lacks accurate V gene annotations and HLA class II information. After controlling the FDR *<* 0.1, we identified 45 positive TCR-HLA pairs for HLA class I.

### TCR generation probability (TGP)

The diversity of TCRs arises from the “V(D)J recombination” process [2]. A TCR *β* chain is generated by randomly selecting and recombining three gene segments, one from each of three types of gene segments: 50 V segments, 2 D segments, and 13 J segments, followed by addition or deletion of nucleotides at the junctions between these segments. A TCR *α* chain is generated by recombination of one of ∼45 V segments and one of 50-60 J segments, with addition or deletion of nucleotides at the segment junctions.

Many groups have attempted to model the V(D)J recombination process [10, 11, 12, 19]. Murugan et al. [11] characterized V(D)J recombination by probabilistic models, parameterizing each event affecting V(D)J recombination according to its interdependencies with other events. Following this work, Zachary et al. utilized dynamic programming to develop OLGA [14], an efficient and flexible algorithm to calculate the theoretical generation probability (TGP), which showed an excellent agreement with published data.

In our study, we aim to quantify the association between an HLA and a TCR using the DePTH score calculated based on their sequences. The TGP of a TCR reflects an underlying property of its sequence, and we seek to evaluate its association with the DePTH scores of the corresponding TCRs. The TCR input for DePTH includes the sequences of the V gene and the CDR3 [9], with the flexibility to allow missing information on the V gene. In the TCGA data [16], the V gene information for a TCR is non-unique or missing for the majority of the TCRs, and also for some TCRs from the Delmonte data [5]. Given this context, the OLGA algorithm [14], which calculates the generation probability of CDR3 sequences in the absence of V gene information, is particularly suited for our purposes. Leveraging the OLGA algorithm, we investigated the relationship between the TGP of CDR3 sequences and their corresponding DePTH scores.

## Results

### Validation of DePTH in additional datasets

In the previous work [9], we trained the DePTH model using the Emerson dataset [6] and evaluated its performance using the Emerson data and a few other relatively small datasets. The results demonstrated that DePTH performed well, surpassing the only previously published model, CLAIRE [7], particularly when evaluated on less common HLAs.

In this work, we evaluated DePTH on the Delmonte [5] and the TCGA [16] datasets. The results demonstrate that DePTH continues to exhibit strong performance on those datasets. The AUC values on the Delmonte dataset for HLA-I and HLA-II were 0.73 and 0.66 (Figure 2), respectively. For the TCGA dataset, which only contains information for HLA-I alleles, the AUC reached 0.8 (Supplementary Figure 1). For comparison, the AUC values on the Emerson dataset for HLA class I and HLA class II were 0.82 and 0.79, respectively. It is important to note that the TCGA dataset lacked V gene information and thus the prediction did not use V genes. When we masked the V gene information in the Emerson dataset, the AUC decreased to 0.73. This means that under the same criterion, DePTH has a better performance on the TCGA dataset than on the Emerson dataset.

**Figure 2:**
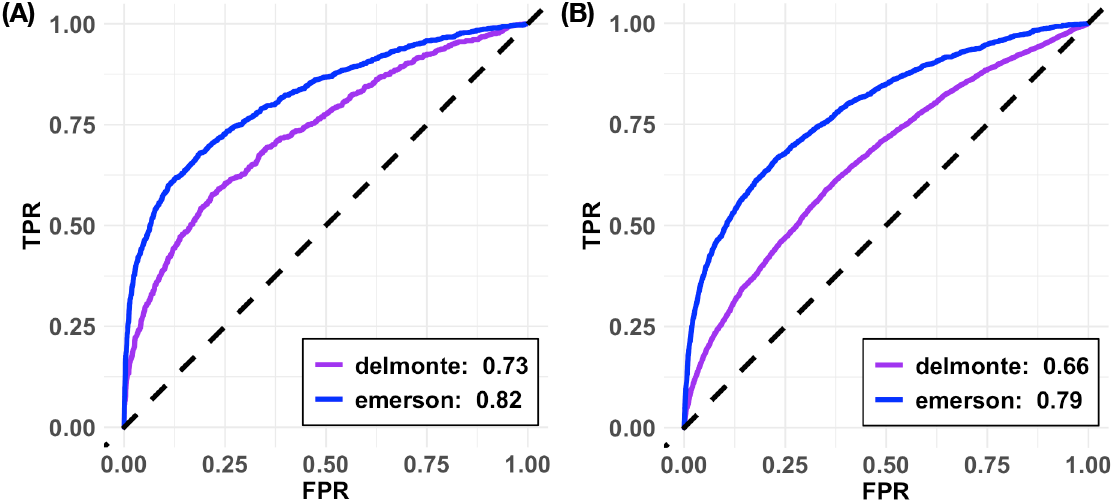
Evaluation of DePTH results on the Delmonte dataset for the associations between TCRs and HLA-I alleles (A) or HLA-II alleles (B).

Beyond the prediction performance of DePTH, it is also important to explore the underlying factors that may distinguish positive (associated) TCR-HLA pairs versus negative (non-associated) ones, and whether DePTH captures information beyond these factors. Towards this end, we mainly focus on two factors: TCR generation probability (TGP) and the length of CDR3 sequence. For the subsequent analyses, we will present the conclusions related to HLA-I from the Delmonte dataset in the main text. The analyses involving HLA-II and the TCGA dataset are included in the Supplementary Materials.

### TCR generation probability (TGP)

We employ the OLGA method [14] to calculate TGPs, which are the probability that a CDR3 is generated by the stochastic V(D)J recombination followed by random insertions/deletions [3].

As expected, the TGP of a TCR is positively correlated with its population frequency (Figure 3(A)). In other words, TCRs with higher TGPs are more likely shared across individuals. TCRs’ population frequency is also an important factor to consider when training DePTH. To prepare the training data for DePTH, we selected the positive TCR-HLA pairs based on their associations (i.e., co-occurrence) in a human population [9], and TCRs with higher population frequencies are more likely to have significant associations. Therefore, without controlling TCR population frequency, the TCRs in positive TCR-HLA pairs have higher TGPs than the TCRs in negative pairs.

**Figure 3:**
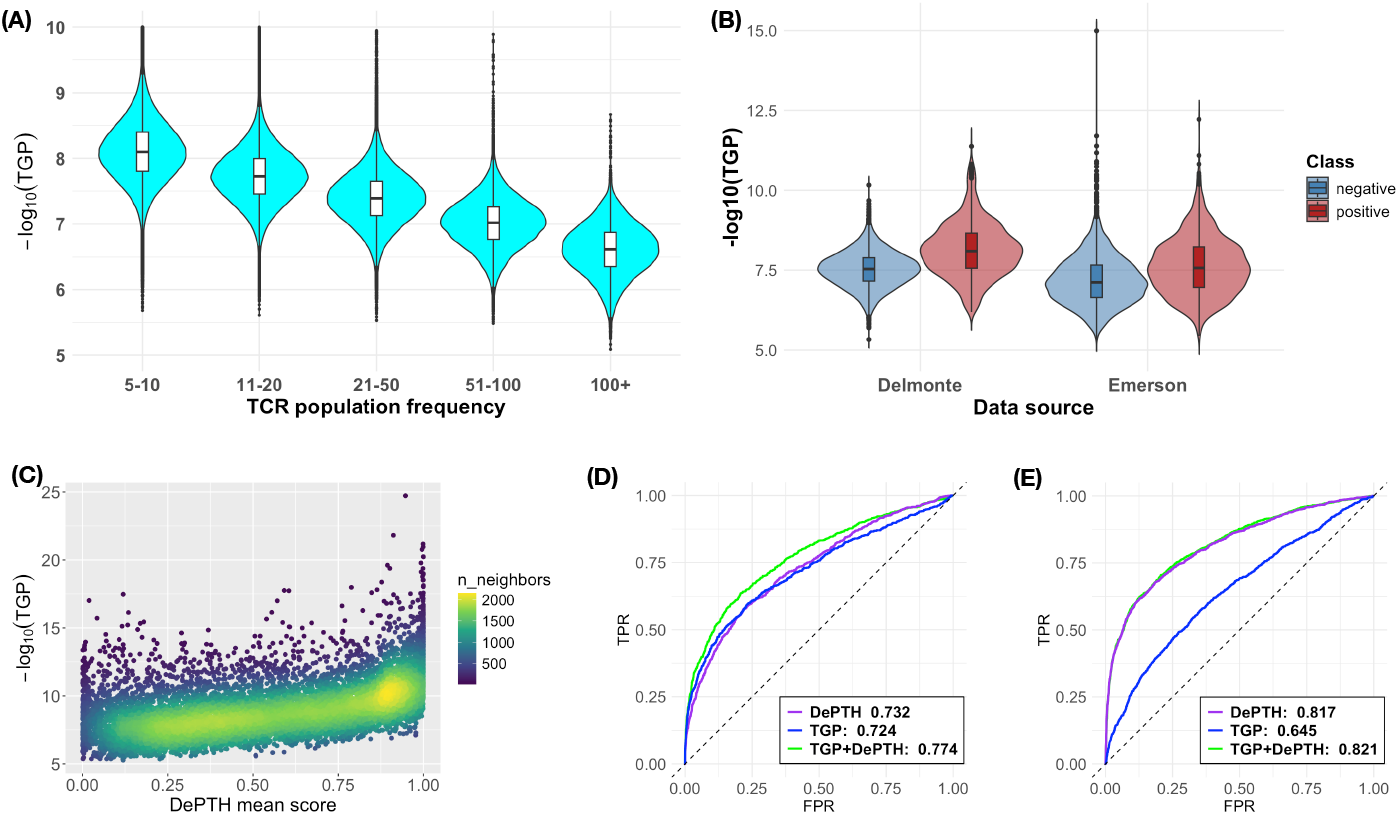
(A) TCR population frequency in Delmonte data versus − log_10_(TGP). (B) The distribution of − log_10_(TGP) of the TCRs that appear in the negative and positive TCR-HLA pairs. (C) DePTH mean score (average across HLAs for each TCR) versus − log_10_(TGP) for a randomly selected patients’ top 10,000 TCRs with highest clonal frequencies. (D) ROC Curves on Delmonte data for TCR-HLA association prediction by three models: DePTH, TGP, and TGP+DePTH. For the combined model, we use logit(*Y*) = *X*_1_ + *X*_2_ + *X*_1_ * *X*_2_ to predict *Y*, the probability of being a positive pair, where *X*_1_ is the DePTH score of the TCR-HLA pair, and *X*_2_ is the TGP of the TCR. (E) The same evaluation as (D) for Emerson dataset.

When we prepared the training data for DePTH [9], we did select the TCRs in the negative TCR-HLA pairs so that their population frequencies matched with those in the positive pairs. However, TGPs are still systematically different between the positive and negative pairs, for the following reasons. Conditioning on a specific population frequency, a TCR from a negative pair may be observed in multiple individuals due to its relatively high TGP. In contrast, the TCRs that appear in the positive pairs may be shared across individuals because they target a common antigen-HLA complex (e.g., a peptide of a common virus presented by a common HLA allele). As a result, after matching the TCR population frequency between positive and negative pairs, the TCRs in positive pairs tend to have lower TGPs than the TCRs in negative pairs (Figure 3(B)). In fact, our DePTH model captures this difference. For each TCR, we define its *DePTH mean score* as the average DePTH score across all HLAs and it is negatively associated with TGP (Figure 3(C)).

Since TGPs of TCRs in positive and negative pairs could be different, we evaluated whether it could be used to distinguish positive vs. negative TCR-HLA pairs. It’s surprising that TGP alone can provide excellent predictions for the Delmonte dataset, achieving an AUC of 0.72, which is close to the performance of DePTH (Figure 3(D)). However, TGP’s performance is strongly influenced by the input data. In the Emerson data, the TGP has much lower prediction AUC than DePTH (Figure 3(E)). The difference of TGP’s performances between Delmonte and Emerson data is likely due to the difference of sample composition. All the individuals in the Delmonte data are patients with acute COVID-19. Thus the positive TCRs of Delmonte data are likely responding to SARS-CoV-2 infection, with lower TGP (Figure 3(E)). This creates stronger contrast with negative TCRs which were selected based on TCR population frequencies. In addition, for the Delmonte dataset, if we don’t match the TCR population frequency between positive and negative pairs, DePTH can still give useful prediction (AUC = 0.67) while TGP’s prediction is much worse (AUC = 0.58).

Next, we explore when TCR population frequency is matched between positive and negative pairs, whether combining TGP and DePTH improves the prediction accuracy. Combining TGP and DePTH does improve the performance for the Delmonte dataset (Figure 3(D)), but it does not make much difference in the Emerson dataset (Figure 3(E)). A plausible explanation is that TGP cannot clearly distinguish positive and negative pairs in the Emerson dataset.

### The impact of CDR3 sequence’s length

In the implementation of DePTH, we padded all CDR3 sequences to 27 amino acids, allowing us to process TCRs of varying lengths together. However, the CDR3 length of a TCR could affect the TCR-HLA interaction, and thus we included CDR3 length before padding as an input to our neural network. We examined the lengths of CDR3 sequences in the Delmonte and Emerson datasets and found that positive TCRs generally had longer CDR3 lengths, regardless of the dataset examined (Figure 4(A-B)). This difference of CDR3’s length distribution supports the incorporating of CDR3 length as an input to our model. In fact, DePTH scores capture information of CDR3 length. TCRs with longer CDR3 sequence tends to have higher mean DePTH score (Figure 4(C)). This observation raises a question that whether DePTH performance varies across CDR3 lengths. In fact, when evaluating DePTH performance across CDR3 lengths, the performances are similar (Figure 4(D)). We further explored whether adjusting training data to match the CDR3 lengths of positive and negative TCRs could help model training. Although the overall AUC of the new length matching model was comparable to the original model (0.73(original) versus 0.72(length matching), respectively), the revised model struggled to consistently deliver accurate predictions across CDR3 lengths (Figure 4(E)). Therefore, we conclude that the default training data, which reflect CDR3 length distribution difference, is a better strategy to prepare training data.

**Figure 4:**
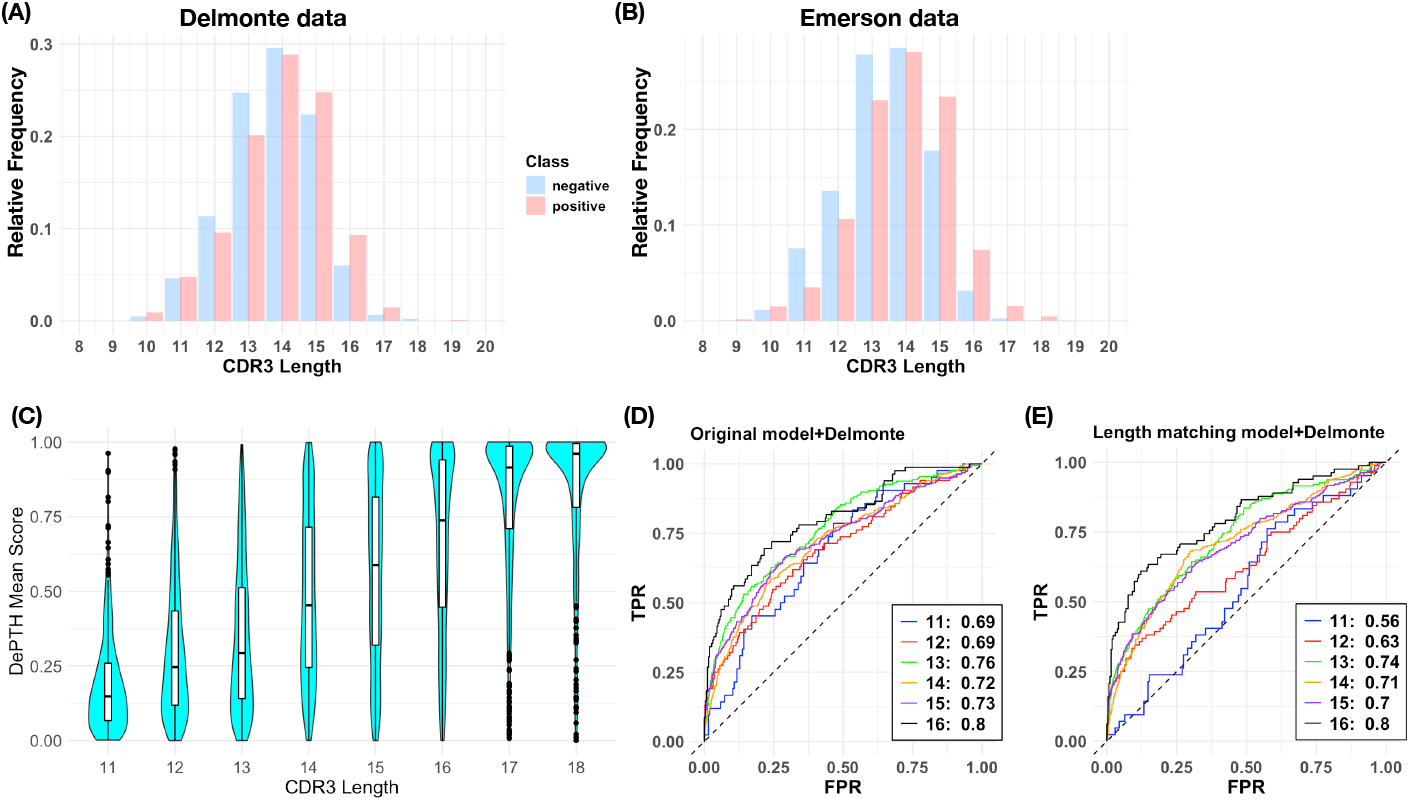
(A-B) The distributions of CDR3 length in Delmonte dataset (A) and Emerson dataset (B). (C) DePTH mean score versus CDR3 length. (D-E) ROC curves evaluating DePTH prediction in Delmonte dataset using default training data and length matching training data, respectively, across CDR3 length.

### DePTH2.0

Since increasing the amount and diversity of training data could improve the performance of the trained model, we are interested in the performance of DePTH when we combine different training datasets. To this end, we compare the performance of three models: the original DePTH trained on Emerson data (DePTH 1.0), DePTH trained on Delmonte data, and DePTH trained on combined Delmonte and Emerson data (DePTH 2.0). We split both Emerson data and Delmonte data to training and test sets with a split ratio of 4:1. Then we evaluated different models in the Emerson or Delmonte test data. DePTH 2.0 gives the best prediction (Figure 5). In particular, DePTH 2.0 outperforms the model trained using Delmonte data, indicating that Emerson data is helpful for learning HLA-TCR associations in the Delmonte dataset. During each training iteration, the Emerson dataset is fixed, while the Delmonte dataset is randomly split into two parts: one for training and one for testing. This random split is conducted five times,tThe results shown in (Figure 5) represent one of the splits and the complete table summarizing all five splits is provided in the supplementary. We attempted multiple random splits of the Delmonte data to train the DePTH model. The results demonstrate that the model, when trained on a combination of Emerson and Delmonte datasets, consistently achieves excellent predictive performance on both datasets.

**Figure 5:**
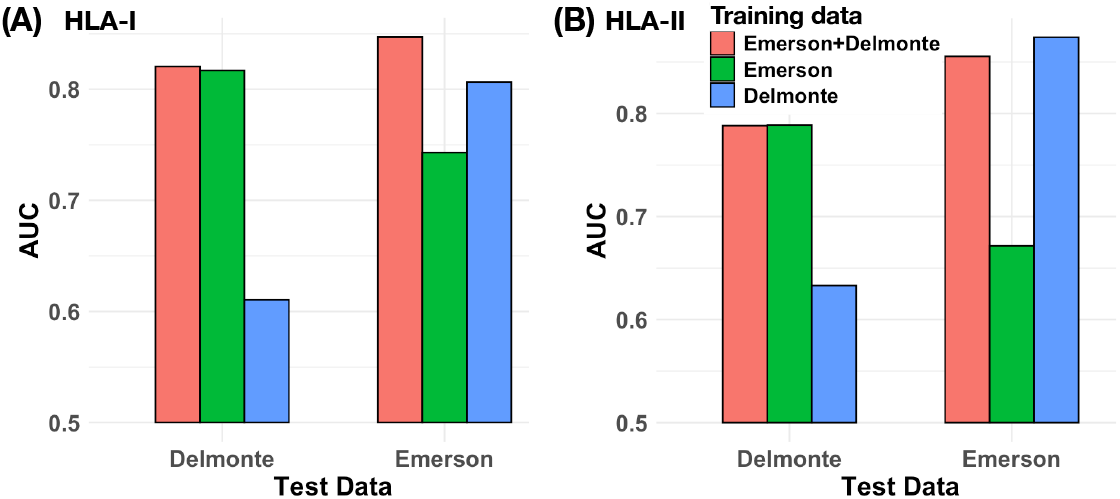
Comparison of the performance of DePTH on Delmonte and Emerson when trained on different datasets (Emerson, Delmonte and Emerson+Delmonte). (A) and (B) represent HLA-I and HLA-II respectively.

## Discussion

In this paper, we performed a systematic evaluation of DePTH. When tested on an independent dataset by Delmonte et al. [5], where TCR were measured by bulk sequencing of TCR beta chains, the DePTH model demonstrated robust prediction on HLA-I (AUC = 0.73). However, the prediction accuracy on HLA-II was lower, with an AUC of 0.66, likely due to the relative complexity of HLA-II. We also evaluated DePTH in a different type of data, the TCGA data [16], where the TCR were extracted from bulk RNA-seq data. The TCGA data only have information of HLA-I alleles, and DePTH can predict the associated TCR-HLA pairs with decent accuracy (AUC = 0.8).

We also studied the role of TCR generation probability (TGP) and CDR3 length in studying the relationship between HLA and TCR. TCRs associated with HLAs (positive TCRs) have lower TGPs than other TCRs (negative TCRs) with similar population frequencies. It is likely that the positive TCRs are popular because they target popular antigens. In contrast, the negative TCRs are popular if they have higher TGPs. We observed a moderate correlation between the TGP of a TCR and the mean DePTH score of this TCR. We also demonstrate that it is possible to combine DePTH score and TGP to improve the accuracy to predict TCR-HLA association, though the performance may depend on the datasets.

We found a moderate correlation between the CDR3 length of a TCR and the mean DePTH score of this TCR. Nevertheless, DePTH performs well for each CDR3 length separately when stratifying the data by CDR3 length. We have also sought to re-train the model using TCRs with matching CDR3 length distributions. However, the length-matching model failed to provide stable predictions across different CDR3 lengths. This suggests that the natural distribution of CDR3 lengths conveys additional information, which should be kept during model training.

Ultimately, we found that incorporating Emerson data into the training set was beneficial for the task of classifying pairs in the Delmonte data. Models trained with both Emerson and Delmonte data consistently outperformed those trained solely with Delmonte data.

We have also noticed that for some TCRs, DePTH’s prediction are consistently high or low regardless of the HLAs, leading to high or low mean-DePTH scores. Both TGP and CDR3 length cannot explain such TCR-specific effects. (Figure 3(C), 4(D)). The top 100 TCRs that consistently have high DePTH scores form a few clusters, when evaluating distances between TCRs using their amino acid sequence similarities. So does the bottom 100 TCRs (Supplementary Figure 5). These results suggest that TCR sequence may play a role here. Addressing this limitation warrants future work, as we aim to enable DePTH to fully integrate both HLA and TCR information, rather than relying primarily on TCR information.

## Code Availability

The code for the analysis can be found at: https://github.com/FUminlee/TCR-project/tree/main.

