## Supplementary for "Improved Deep Learning Prediction of TCR-HLA Associations"

### DePTH2 Supplementary

November 23, 2024

#### Prepare HLA-TCR pairs

The dataset we used to evaluate DePTH [3] included the Delmonte [2] and the TCGA [4] datasets. We only considered the HLA and TCRs with population frequency equal to or greater than 5 for either dataset. Each TCR was composed of a CDR3 sequence and a V gene. When a TCR record in the dataset corresponded to multiple V genes, we retained the first one. For TCGA dataset, we excluded the V gene information because most TCRs either had missing V gene information or were associated with more than one V genes.

##### Positive HLA-TCR pairs

We selected positive HLA-TCR pairs (i.e., the HLA-TCR pairs that tended to co-occur in a population of samples) by Fisher's exact test. First, the Fisher's exact p-value for a HLA-TCR pair was calculated based on their co-occurrence in the population. Next, we selected positive pairs while controlling the FDR to be less than 10%. We leveraged a conservative method to calculate FDR by assuming that all HLA-TCR pairs were negative. For a p-value threshold $\alpha$, the expected number of pairs that had p-values smaller than $\alpha$ by chance was $M \times \alpha$, where $M$ was the total number of HLA-TCR pairs that we considered. We chose the p-value threshold $\alpha_1 = \arg\max_\alpha \frac{M \times \alpha}{m_\alpha} < 10\%$, where $m_\alpha$ was the total number of pairs whose p-values were less than $\alpha$.

##### Negative HLA-TCR pairs

When selecting negative HLA-TCR pairs, we maintained a ratio of five negative pairs versus one positive pair. In addition, we maintained the marginal distribution of HLA alleles with 5:1 ratio. For instance, if a particular HLA allele appeared twice in the positive pairs, we ensured it was represented ten times in the negative pairs. Furthermore, for the Delmonte dataset, we matched the distribution of TCR population frequencies between the negative and positive pairs. For the TCGA dataset, due to its sparsity, we were unable to match the TCR population frequency distribution of positive and negative pairs.

### TCGA analysis

The TCGA dataset is different from the Delmonte and Emerson datasets in three aspects. First, its TCR data were derived from RNA-seq data and thus each sample only had a small number of TCRs. Secondly, it contained only HLA-I information for participants. Thirdly, it lacked accurate V gene information for most of the TCRs. Therefore, our input data is restricted to HLA-I alleles and CDR3 sequences for TCRs. Despite this limitation, the DePTH model demonstrated remarkable performance on the TCGA dataset, achieving an AUC of 0.8 (Figure 1(A)). In contrast, when V gene information was excluded from the test data derived from Emerson data, the model's performance was slightly worse, with an AUC of 0.73.

#### TCR generation probability

In the TCGA dataset, the TGPs of the negative TCRs (the TCRs from the negative pairs) were higher than those of the positive TCRs (the TCRs from the positive pairs), and this trend was more pronounced in the TCGA data compared to Delmonte or Emerson (Figure 1(B)). Additionally, both positive and negative TCRs in the TCGA dataset exhibited higher TGPs than those in Delmonte or Emerson data. This was likely due to the sparsity of the TCGA data. More specifically, when only tens or hundreds of TCRs were sampled from one sample, those TCRs were more likely to have higher TGPs. The TGP and TCR population frequency were positively correlated, which was consistent with the observations from Delmonte or Emerson data (Figure 1(C)). TGP effectively distinguishes between positive and negative pairs within the TCGA dataset. However, the DePTH model captures additional information. When we combined the TGP and the DePTH score using a linear model, the accuracy of the model improved (Figure 1(D)).

#### CDR3 Length Analysis

We investigated the length distributions of positive and negative TCRs and compared the performance of the original DePTH model and the Length Matching model across different TCR lengths in the TCGA dataset. The length distribution of positive and negative TCRs exhibits striking differences (Figure 2(A)). Positive TCRs are predominantly between 13 and 14 amino acids in length, whereas negative TCRs are more commonly found within the 11-13 length range. This trend, where positive TCRs tend to be longer than negative TCRs, is also observed in the Delmonte and Emerson datasets. However, the distinction between the length distributions of positive and negative TCRs is even more pronounced in the TCGA dataset. In the TCGA data, both the original DePTH model and the Length Matching model have highly variable performances across CDR3 lengths (Figure 2(B-C)), which may be partly due to the small number of TCRs for each CDR3 length.

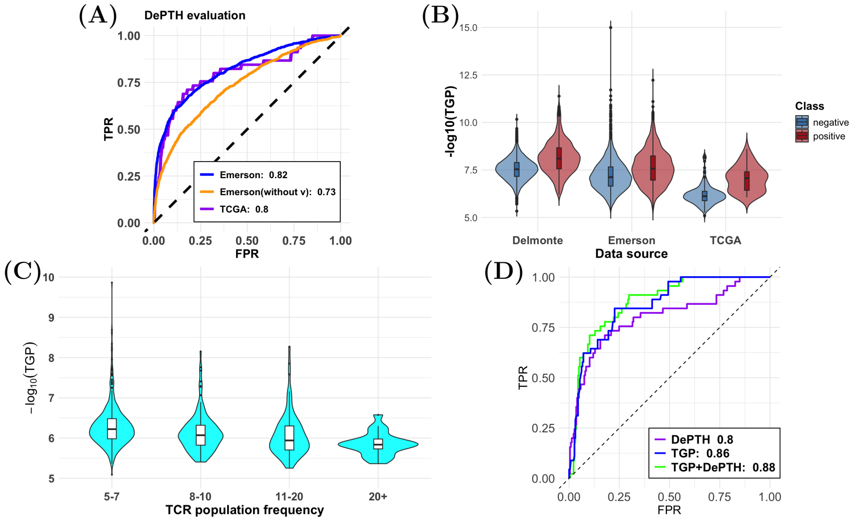


Figure 1: (A) ROC curves comparing the performance of the DePTH model on the TCGA and Emerson datasets. The legend shows the AUCs for the Emerson dataset with (AUC = 0.82) or without V gene (AUC = 0.73), and the AUC for the TCGA dataset (AUC = 0.8). (B) Violin plots illustrating the distribution of $-\log_{10}(\text{TGP})$ for positive (red) and negative (blue) TCRs across the Delmonte, Emerson, and TCGA datasets. (C) The distribution of $-\log_{10}(\text{TGP})$ for different TCR population frequencies on TCGA dataset. (D) The ROC curves for three models: DePTH, TGP, and TGP+DePTH on TCGA dataset. The x-axis represents the false positive rate (FPR), and the y-axis represents the true positive rate (TPR). The AUC of DePTH, TGP, and TGP+DePTH is 0.8, 0.86, and 0.88, respectively.

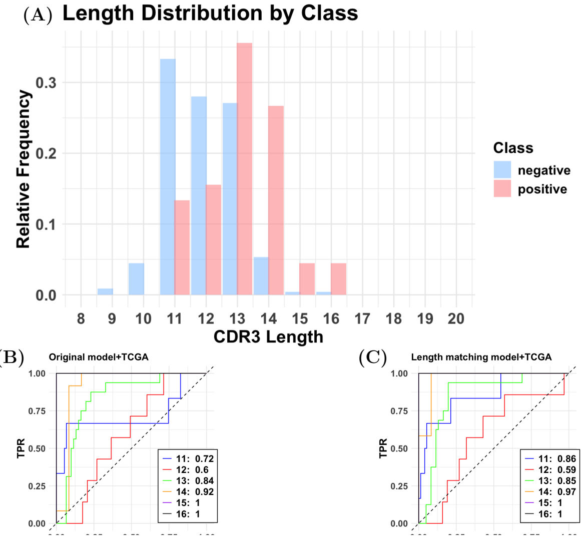


Figure 2: (A) Length distribution of TCRs in the TCGA dataset, divided into positive (red) and negative (blue) classes. (B-C) ROC curves for the original DePTH model (B) and TCR CDR3 length matching model (C), applied to the TCGA dataset, with AUC values for lengths 11 to 16 are shown in the legend.

### Delmonte HLA-II analysis

#### TCR generation probability(TGP)

Similar to the HLA-I analysis, we compared the TGPs between positive and negative TCRs, evaluated the relation between TGPs and DePTH mean scores, and investigated the role of TGP in classifying positive and negative pairs.

In both the Delmonte and Emerson datasets, positive TCRs tend to have lower TGPs than negative TCRs (Figure 3(A)), which is consistent with the observations for HLA-I analysis. Next, we investigated the relationship between the DePTH mean scores of TCRs and their TGPs. The DePTH mean score of a TCR is the mean value of the DePTH scores between this TCR and all HLA-II alleles included in our analysis. We selected the top 10,000 TCRs from one participant and analyzed the relationship between TGPs and DePTH mean scores. The results revealed a clear negative correlation between TGPs and DePTH mean scores (Figure 3(B)). Finally, we evaluated whether TGP could classify positive versus negative TCR-HLA pairs. We separately analyzed the classification performance of TGP on the Delmonte and Emerson datasets. The results show that TGP demonstrates high accuracy in distinguishing positive pairs from negative pairs in the Delmonte dataset (Figure 3(C)). However, its accuracy decreases when applied to the Emerson dataset (Figure 3(D)).

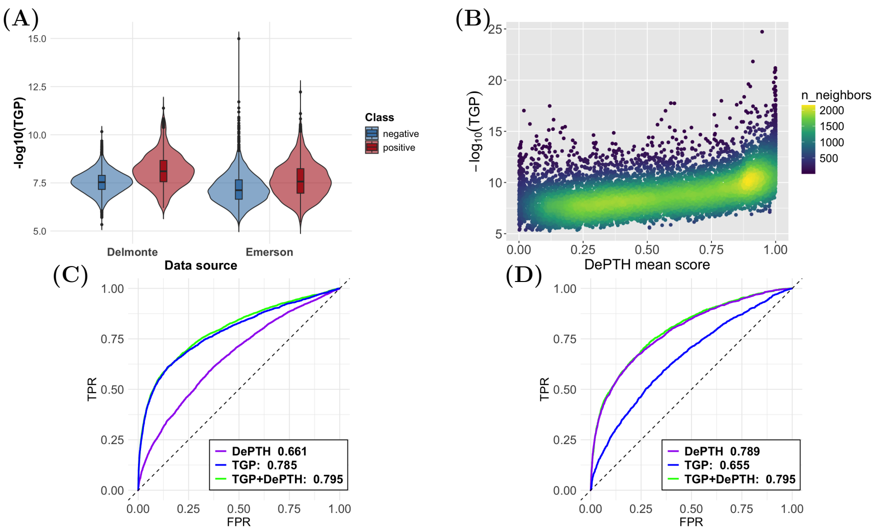


Figure 3: Exploration of HLA-II: (A) The distribution of the TGPs of the TCRs that appear in the negative and positive HLA-TCR pairs. (B) $-\log_{10}(TGP)$ versus mean DePTH score for a randomly selected patient' top 10000 TCRs for Delmonte dataset. (C) ROC curves on Delmonte for different models: DePTH, TGP, and TGP+DePTH. Here in the case of TGP+DePTH, we use the model $\text{logit}(Y) = X_1 + X_2 + X_1 * X_2$ to predict Y, where $X_1$ is the DePTH score, $X_2$ is the TGP and $Y$ is the probability of belonging to positive set. (D) The same evaluation as (C) for Emerson dataset.

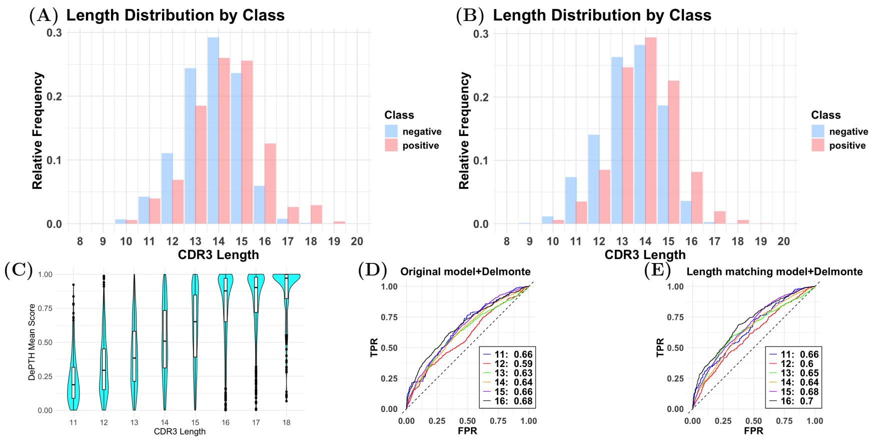


Figure 4: (A-B) The distribution of the CDR3 lengths for the TCRs in the negative and positive TCR-HLA-II pairs for Delmonte dataset (A) and Emerson dataset (B). (C) The distribution of DePTH mean scores for TCRs with different CDR3 lengths, ranging from 11 to 18. The boxplots inside the violin plots indicate the median and quartiles of each distribution. (D-E) ROC curves evaluating DePTH prediction in Delmonte dataset using either default training data (D) or length matching training data (E) across CDR3 length.

#### CDR3 length analysis

We compared CDR3 lengths between TCRs from positive or negative pairs, investigated the relationship between CDR3 length and the DePTH mean score, and evaluated the performance of the original DePTH model and the Length Matching model across different TCR lengths.

The CDR3 of negative TCRs tend to be shorter than those of positive TCRs (Figure 4(A-B)), which is consistent with the findings for HLA-I analysis. TCRs with longer CDR3s tend to have larger DePTH mean scores (Figure 4(C)), which is also consistent with observation for HLA-I. The original DePTH model and the Length Matching DePTH model both have similar performances across CDR3 lengths(Figure 4(D-E)). In contrast, for HLA-I data, the original DePTH model has robust performance across CDR3 lengths, but the Length Matching model's performance is more variable across CDR3 lengths.

#### TCRs with high or low DePTH mean scores

Some TCRs have DePTH mean scores (i.e., the average of the DePTH scores between this TCR and all the HLAs) around 0 or 1. This implies that altering HLA information results in limited variance in the DePTH scores for these TCRs.

We examined whether the TCRs with extreme DePTH mean scores tended to

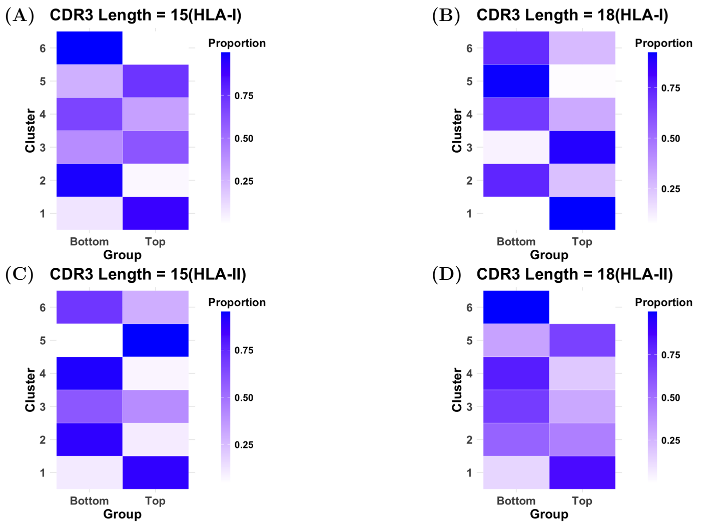


Figure 5: (A) For TCRs with CDR3 Length = 15, the proportion of TCR sequences in the Top set and Bottom set, where the Top and Bottom set were divided based on the DePTH mean score under HLA-I. The Leiden clustering was applied to group the TCRs, and the heatmaps display the distribution of TCRs in each cluster between the Top and Bottom sets. The color intensity indicates the proportion of TCRs in the Top and Bottom sets, with darker colors representing higher proportions. (B) The same case for TCR sequences with CDR3 Length = 18. (C-D) The same case as (A-B) under HLA-II.

have similar sequences. For each CDR3 length, we selected 100 TCRs from the Top set (those with highest DePTH mean scores) and 100 from the Bottom set (those with lowest DePTH mean scores). We calculated the distance between every two of the 200 TCRs using tcrdist [1]. Then, we applied a distance cutoff to create a TCR-TCR graph to be used for Leiden clustering. The majority of clusters are predominantly composed of TCRs from either the Top set or the Bottom set. These results show that the two groups of TCRs from the top set and bottom set have different sequences, and within each group, the TCRs form some clusters based on their sequence similarities. Therefore, the sequences of the TCRs can be informative to separate those TCRs with extreme DePTH scores, while there are still heterogeneity of TCR sequences within top or bottom set (Figure 5).

### Combined training full table

During the combined training process, we conducted five experiments in which the Delmonte data was randomly split into training, validation, and test sets.

Table 1: Model AUC Comparison(HLA-I)

| Training | Test | Split1 | Split2 | Split3 | Split4 | Split5 |
|---|---|---|---|---|---|---|
| Emerson | Emerson | 0.82 | 0.82 | 0.82 | 0.82 | 0.82 |
| Delmonte | Emerson | 0.51 | 0.60 | 0.61 | 0.61 | 0.61 |
| Emerson+Delmonte | Emerson | 0.82 | 0.82 | 0.82 | 0.82 | 0.82 |
| Emerson | Delmonte | 0.75 | 0.72 | 0.74 | 0.73 | 0.71 |
| Delmonte | Delmonte | 0.78 | 0.82 | 0.81 | 0.81 | 0.82 |
| Emerson+Delmonte | Delmonte | 0.83 | 0.82 | 0.85 | 0.84 | 0.81 |
| Emerson | TCGA | 0.80 | 0.80 | 0.80 | 0.80 | 0.80 |
| Delmonte | TCGA | 0.66 | 0.65 | 0.64 | 0.59 | 0.59 |
| Emerson+Delmonte | TCGA | 0.63 | 0.72 | 0.67 | 0.59 | 0.65 |

Table 2: Model AUC Comparison(HLA-II)

| Training | Test | Split1 | Split2 | Split3 | Split4 | Split5 |
|---|---|---|---|---|---|---|
| Emerson | Emerson | 0.79 | 0.79 | 0.79 | 0.79 | 0.79 |
| Delmonte | Emerson | 0.63 | 0.63 | 0.63 | 0.63 | 0.63 |
| Emerson+Delmonte | Emerson | 0.79 | 0.79 | 0.79 | 0.79 | 0.79 |
| Emerson | Delmonte | 0.67 | 0.66 | 0.66 | 0.66 | 0.67 |
| Delmonte | Delmonte | 0.87 | 0.89 | 0.89 | 0.88 | 0.88 |
| Emerson+Delmonte | Delmonte | 0.86 | 0.87 | 0.86 | 0.86 | 0.86 |

Each time, the combined model demonstrated good performance, maintaining high accuracy on the Emerson, Delmonte, and TCGA data. (Table 1 and 2)

#### Read-depth

We explored the read-depth characteristics of Delmonte and Emerson. Our comparison focused on two key aspects: the number of unique TCRs identified for each participant and the total number of TCR templates. The results indicate that the Emerson dataset offers a deeper sequencing depth compared to the Delmonte dataset, under both criteria (Figure 6).

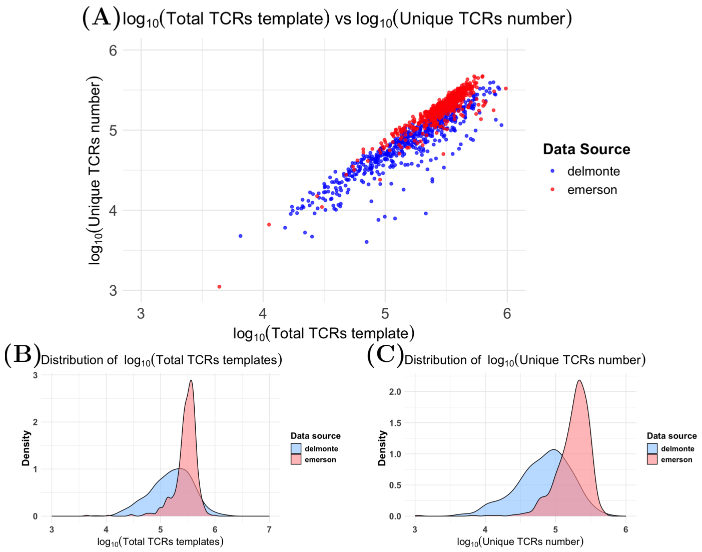


Figure 6: (A) The scatter plot shows the relationship between the total TCR templates and the unique TCRs number on a log scale. Each point represents a participant.(B) Density plot of the log-transformed total TCR templates for each dataset. (C) Density plot of the log-transformed unique TCRs number for each dataset.
